# Generation of scale-invariant sequential activity in linear recurrent networks

**DOI:** 10.1101/580522

**Authors:** Yue Liu, Marc W. Howard

## Abstract

Sequential neural activity has been observed in many parts of the brain and has been proposed as a neural mechanism for memory. The natural world expresses temporal relationships at a wide range of scales. Because we cannot know the relevant scales *a priori* it is desirable that memory, and thus the generated sequences, are scale-invariant. Although recurrent neural network models have been proposed as a mechanism for generating sequences, the requirements for *scale-invariant* sequences are not known. This paper reports the constraints that enable a linear recurrent neural network model to generate scale-invariant sequential activity. A straightforward eigendecomposition analysis results in two independent conditions that are required for scale-invariance for connectivity matrices with real, distinct eigenvalues. First, the eigenvalues of the network must be geometrically spaced. Second, the eigenvectors must be related to one another *via* translation. These constraints are easily generalizable for matrices that have complex and distinct eigenvalues. Analogous albeit less compact constraints hold for matrices with degenerate eigenvalues. These constraints, along with considerations on initial conditions, provide a general recipe to build linear recurrent neural networks that support scale-invariant sequential activity.

## 1 Introduction

The current state of the brain can carry memory for the past *via* history-dependent dynamics. This memory can be used to adaptively shape behavior to anticipate the future. However, the natural world has temporal relationships on a wide range of timescales (e.g. Voss and Clarke, 1975). This presents a problem in the design of the history-dependent dynamics of the brain. The world can contain behaviorally relevant predictive information over a range of scales, but we do not necessarily know the relevant timescales *a priori.* Suppose the brain’s dynamics had a single characteristic scale *s_o_*. If the world contains useful information at a scale much longer than *s_o_*, this information would be invisible to the brain’s dynamics and the system would not be able to exploit this information. Similarly, if the world contains useful information at a scale much shorter than *s_o_*, the brain’s dynamics would not represent this information efficiently. One solution is a dynamic representation of the world that is scale-invariant across time (Howard and Shankar, 2018). Indeed, a spectrum of timescales is an essential ingredient in neural circuit models for temporal pattern recognition (Tank and Hopfield, 1987; Hopfield and Brody, 2000; Buonomano and Maass, 2009; Gütig and Sompolinsky, 2009).

There is empirical evidence suggesting that the brain in fact implements something like scale-invariance. Decades of research in cognitive psychology demonstrate that human timing and memory behavior exhibit the same properties on a wide range of timescales (Murdock, 1962; Glenberg et al., 1980; Rakitin et al., 1998; Howard et al., 2008). Sequential neural activity has been observed in many areas of the brain and is thought to have important cognitive functions in memory and decision making (MacDonald et al., 2011; Harvey et al., 2012; Howard, 2018). In light of these considerations, recently it has been proposed that “scale-invariance” is a desirable property for neural sequences (Shankar and Howard, 2012, 2013). In a scale-invariant neural sequence, the cells that are activated later have wider temporal receptive fields. More specifically, the responses of different cells have identical time courses when they are rescaled in time by their peaks. The hypothesis of scale-invariant neural sequences for time is consistent with recent electrophysiological recordings of “time cells” during a delay period when animals are performing various cognitive tasks (Pastalkova et al., 2008; Jin et al., 2009; MacDonald et al., 2011; Kraus et al., 2013; Mello et al., 2015; Salz et al., 2016; Tiganj et al., 2018). The firing fields of time cells that fire later in the delay period are wider than the firing fields of time cells that fire earlier in the delay period.

Many researchers have studied recurrent neural networks that generate sequential activity (Goldman, 2009; Rajan et al., 2016; Wang et al., 2018), but not many of these works considered scale-invariant sequences (but see Voelker and Eliasmith, 2018). In this paper we seek to identify general constraints on the network connectivity for the generation of scale-invariant neural sequences in recurrent neural networks. We study a linear network of interacting neurons. The f-I curve of many neurons are observed to be largely linear (e.g. Chance et al., 2002). The learning dynamics of linear feedforward neural networks exhibit many similarities compared to their non-linear counterparts (Saxe et al., 2013). Therefore linear neural networks provide a good model for studying systems-level properties of real neuronal circuits.

The paper is organized as follows. In Section 2 the network constraints for the generation of scale-invariant neural sequences are derived analytically. The exact constraints hold when the connectivity matrix has real, distinct eigenvalues. The modifications to generalize them to complex and distinct eigenvalues are discussed. Networks with degenerate eigenvalues are discussed in Appendices A.3 and A.4. To illustrate the mathematical result, in Section 3 two example networks with different single cell dynamics are constructed. Each of the constraints are broken to show that they are necessary for scale-invariance. In Section 4, we compare the eigenvectors and eigenvalues of a chaining model and a random recurrent network with those of the examples in Section 3. It is shown that neither of these networks satisfy the structural constraints derived from Section 2, therefore neither of them support sequential neural activity that is scale-invariant.

## 2 Derivation of the constraints for scale-invariance

In this section we derive the constraints on the connectivity matrix of a linear recurrent network for it to support scale-invariant activity. In Section 2.1, we start with a formal definition of scale-invariance of network activity. In Section 2.2 and Section 2.3 we will derive the two constraints on the connectivity matrix to achieve scale-invariance. Lastly in Section 2.4, we point out that in addition to the two constraints, a particular initial condition is required for the subsequent network dynamics to be scale-invariant. These constraints are sufficient and necessary for a network to generate scale-invariant activity, but only if the network connectivity matrix has real, distinct eigenvalues. We mention a straightforward modification to the results when the network has complex but distinct eigenvalues. The details are shown in Appendix A.1. We discuss the modifications to the constraints for matrices with degenerate eigenvalues in Appendices A.3 and A.4. We also show in Appendix A.5 that deviations from the constraints derived below will cause a graceful degradation in the scale-invariant property of the resulting network activity.

### 2.1 Formulation of the problem

We consider the autonomous dynamics of a linear recurrent network with *N* neurons:

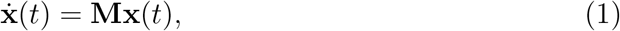

where **x** is an *N*-dimensional vector summarizing the activity of all the neurons in the network and **M** is the *N* × *N* connectivity matrix of the network. We consider the case where there is no input into the network since sequential neural activity is thought to be maintained by internal neuronal dynamics (Pastalkova et al., 2008).

Scale-invariance of the network activity means that the responses of any two neurons in the sequence are rescaled version of each other in time (Figure 1b). Mathematically, this requirement can be written in the following form:

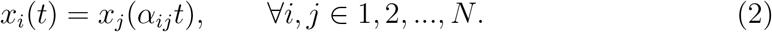

**Figure 1:**
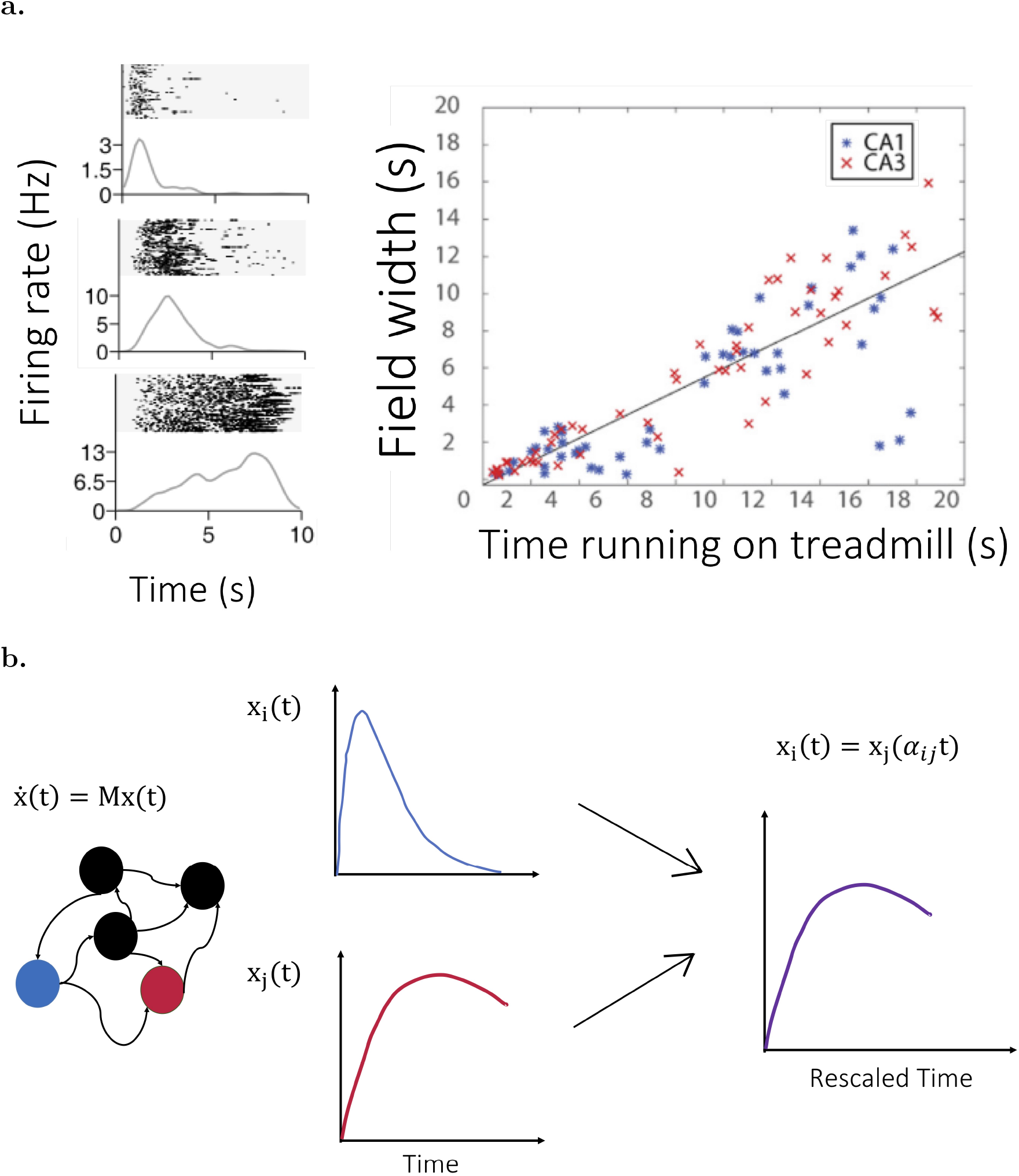
Scale-invariance for neural sequences and the setup of the problem. **a** Left: the raster plots (top) and trial-averaged firing rates (bottom) for three neurons from MacDonald et al. (2011). The neurons were recorded in the hippocampus of rats during a delay period when they were waiting to sample an odor. The neurons that fire later in the delay period show wider responses than neurons that fire earlier during the delay. Right: a scatter plot showing the relationship between the width of each neuron’s response and the peak time at which that neuron fires for all neurons recorded from Salz et al. (2016)). Neurons were recorded in the CA3 region of the rat hippocampus during the delay period of a T-maze alteration task when the animal was running on a treadmill. **b.** In this work, we study the dynamics of a linear recurrent network. We seek constraints on the network connectivity matrix **M** (**b**, left) such that the activity of every pair of neurons (here for example neurons *i* and *j*, middle) are rescaled version of each other in time (right).

That is, for every pair of neurons *i* and *j*, their responses *x_i_*(*t*) and *X_j_*(*t*) are rescaled in time by a factor *α_ij_*. As will be discussed below, this condition can only be satisfied when the network connectivity has real, distinct eigenvalues. Otherwise, only a subset of responses can be rescaled versions of each other. Therefore, in the following sections we are going to focus on connectivity matrices with real, distinct eigenvalues. We will derive two conditions on the connectivity matrix **M** necessary for Equation 2 to hold and for the network to generate scale-invariant sequential activity. When the connectivity matrix has complex, distinct eigenvalues, we will state the constraints that allow it to have two distinct scale-invariant sequences. The modifications to the constraints when the eigenvalues are degenerate are discussed in Appendices A.3 and Appendices A.4.

### 2.2 Constraint 1: Geometrically spaced network timescales

We start by solving Equation 1 using the standard eigendecomposition technique. We diagonalize the connectivity matrix **M** as **M** = **UΛU**^-1^ where **Λ** is a diagonal matrix consisting of the eigenvalues and **U** is a matrix whose columns are the eigenvectors of **M**. The solution of Equation 1 is then a linear combination of exponential functions:

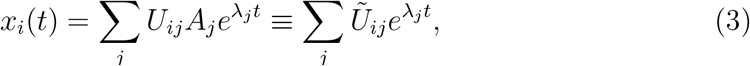

where the *A_i_*’s are constants determined by the initial condition and are absorbed into the definition of the matrix **Ũ**.

Imposing the scale-invariance condition (Equation 2) on Equation 3, we have

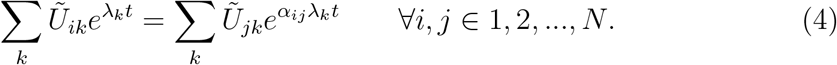

For this equation to hold, the time-dependent parts on both sides of the equation must be identical. This means that all the eigenvalues involved in the left hand side of the equation should be equal to a scaled version of the eigenvalues involved in the right hand side. In other words, for each λ_*k*_ there should exist an integer *δ* such that *λ_k+δ_* = *α_ij_*λ_*k*_. The only way to achieve this is to have a geometric progression of eigenvalues (network timescales) (Figure 2a). For example, λ_1_ = −1, λ_2_ = −2, λ_3_ = −4, etc.. Therefore we arrive at the first constraint (as depicted in Figure 2a):

**Figure 2:**
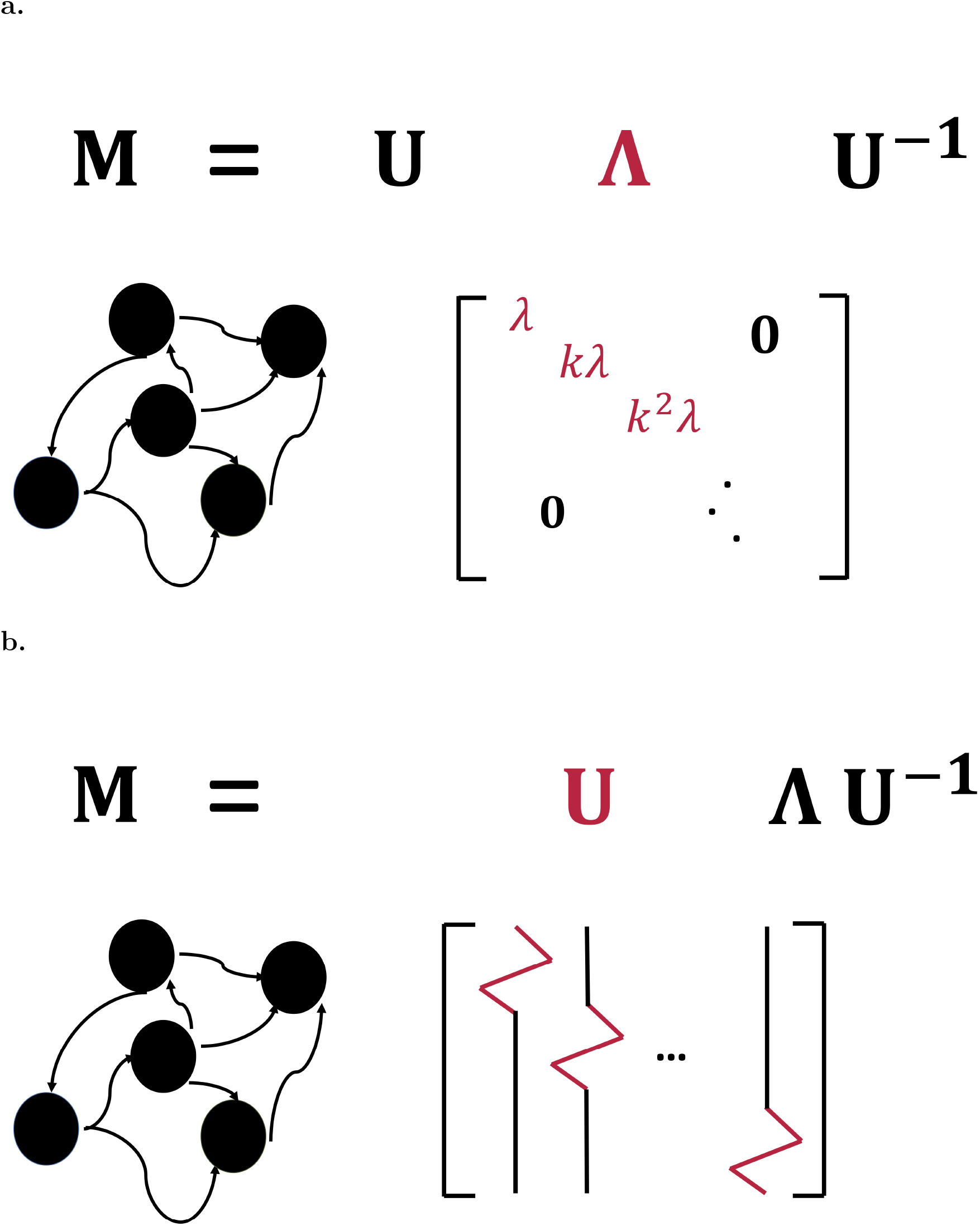
Graphical illustrations of the constraints for scale-invariance in linear recurrent networks whose connectivity matrices are diagonalizable and have real, distinct eigenvalues. To generate scale-invariant activity in linear recurrent networks whose connectivity matrices are diagonalizable and have real and distinct eigenvalues (other cases are discussed in Appendix A.1–A.4), the network connectivity must have geometrically spaced eigenvalues (**a**). Furthermore, it must have translation invariant eigenvectors (columns of the matrix **U**) that consist of the same motif (in red) at translated entries (**b**).

#### Constraint 1

For a linear recurrent network whose connectivity matrix is diagonalizable and has real, distinct eigenvalues to generate scale-invariant activity, the eigenvalues must form a geometric progression.

#### Remarks

The above constraint needs to be slightly modified when there are complex eigenvalues, since the eigenvalues must come in complex conjugate pairs for the connectivity matrix **M** to be real. It can be shown (for details see Appendix A.1) that contrary to the definition of scale-invariance above (Equation 2), there can at best be two scale-invariant sequences, where the responses of neurons in different sequences are not rescaled versions of each other. For this to happen, all the eigenvalues must be complex^1^. Therefore Constraint 1 in the case of complex eigenvalues is as follows (See Appendix A.1 for details):

#### Constraint 1 for connectivity matrices with complex eigenvalues

For a linear recurrent network whose connectivity matrix is diagonalizable and has complex eigenvalues to generate two sequences of scale-invariant activity, the eigenvalues must all be complex and form two geometric progressions that are complex conjugate with each other.

In the example networks constructed in this paper, all of the eigenvalues have negative real parts to prevent unbounded growth of network activity.

### 2.3 Constraint 2: Translation-invariant eigenvectors

A second constraint for Equation 4 to hold is that the rows of **Ũ** must satisfy a translation-invariant relationship. *Ũ_i,k+δ_* = *Ũ_jk_*. This way the modes that different neurons pick out will be rescaled versions of each other. Recall from Equation 3 that *Ũ_ij_* = *Ũ_ij_A_j_*. Therefore the condition above is equivalent to the columns of the matrix **U** being translation-invariant up to a constant. For example, the different columns could be **v**_1_ = [1, −1,0, 0, 0], **v**_2_ = [0,1, −1, 0,0], **v**_3_ = [0, 0,1, −1, 0], etc.. Notice that the columns of **U** are just the eigenvectors of **M**. Therefore, we reach the second constraint (see Figure 2b for a graphical illustration):

#### Constraint 2

For a linear recurrent network whose connectivity matrix has real, distinct eigenvalues to generate scale-invariant activity, the eigenvectors must consist of the same motif (up to a scaling factor) at translated entries. In other words, the eigenvectors must be translation-invariant.

#### Remarks

When the connectivity matrix has complex eigenvectors, they must come in complex conjugate pairs to ensure that the matrix is real. In this case, it can be shown (for details see Appendix A.1) that there can at best be two scale-invariant sequences, contrary to the definition of scale-invariance above (Equation 2). The responses of neurons in different sequences are not rescaled versions of each other. For this to happen, all the eigenvalues must be complex, and Constraint 2 in the case of complex eigenvalues is as follows (see Appendix A.1 for details):

#### Constraint 2 for connectivity matrices with complex eigenvalues

For a linear recurrent network whose connectivity is diagonalizable and whose eigenvalues are complex to generate two sequences of scale-invariant activity, each eigenvector should either be a translated version of another eigenvector or its complex conjugate.

A motif with length *L* in a vector of length N can at most be translated *N – L* times. Therefore, there will be L eigenvectors that are not translated version of the rest of the eigenvectors. In this work we are interested in large networks where *N ≫ L*. Therefore this number is negligible compared to the total number of eigenvectors.

### 2.4 A note on initial conditions

Besides the constraints on the connectivity matrix, the initial condition of the network also affects the scale-invariance of the network activity. Equation 3 requires each neuron in the scale-invariant sequence to have a specific initial condition, i.e. 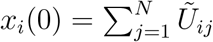, to ensure that the dynamics that ensues are scale-invariant. This holds regardless of the eigenspectrum of the connectivity matrix.

For example, if the network has real, distinct eigenvalues, and the sum of the elements in the motif is 0 (which will be the case for the examples in Section 3), the constraint on the initial condition becomes:

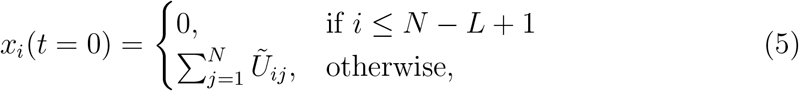

where *L* is the length of the repeating motif. In the case where the number of neurons in the network is much larger than the length of motif (*N≫L*), this constraint on initial condition states that most of the neurons in the network need to be inactive at *t* = 0, whereas a few active neurons act as “input nodes” and propagate their activity to the rest of the network.

## 3 Examples

In this section we construct two example networks based on the analytical results derived above and show that they allow scale-invariant sequential dynamics. The connectivity matrices of these networks will have geometric progressions of eigenvalues and tranlation-invariant eigenvectors, as shown in Section 2. In the first example (Section 3.1), all eigenvalues are real and the neurons in the network have simple unimodal temporal receptive fields. In the second example (Section 3.2), all eigenvalues are complex, which gives rise to more complicated damped oscillatory single neuron dynamics. In this case, there will be two scale-invariant sequences, as mentioned in Section 2. We will also show that the network activity is no longer scale-invariant when either of the two constraints is violated. In Section 3.3, we will discuss the relationship of our results to a previously proposed network model that generates scale-invariant sequential activity (Shankar and Howard, 2013). In what follows, all simulations were performed in Python 3.6 using Euler’s method.

### 3.1 Simple unimodal temporal receptive fields

In this example we constructed a network that generates sequentially-activated cells with a scale-invariant property. The network consists of *N* = 10 cells. Each cell will have a simple unimodal temporal receptive field, similar to what was observed in electrophysiological recordings of “time cells” (Pastalkova et al., 2008; Jin et al., 2009; MacDonald et al., 2011; Kraus et al., 2013; Mello et al., 2015; Salz et al., 2016; Tiganj et al., 2018).

We constructed the connectivity matrix from its eigendecomposition **M** = **UΛU**^-1^. According to Constraint 1 (Section 2.2), the network must have geometrically spaced eigenvalues. We hence let **Λ** be a diagonal matrix whose diagonal elements are geometrically spaced between −0.1 and −5.12.

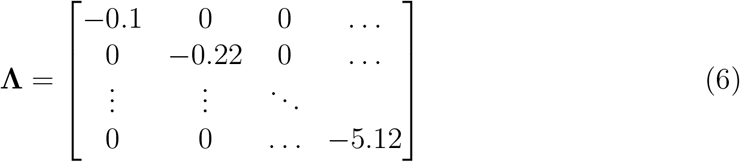

According to Constraint 2, the eigenvectors of the connectivity matrix **M** must consist of the same motif. Equivalently, we constructed the matrix **U** such that its *rows* consist of the same motif. In this example we constructed the motif to be (1, −1). Therefore the matrix **U** was given by

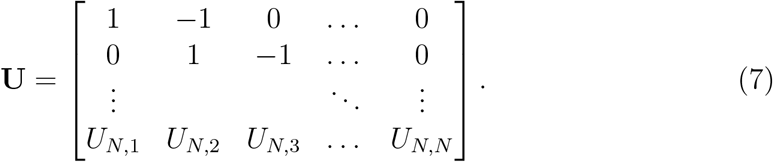

As mentioned in Section 2.3, since the motif in the eigenvectors can at most be translated *N* – 2 times, the *N*th neuron will not be part of the scale-invariant sequence. Therefore, we chose the last row in **U** to be arbitrary numbers such that **U** is invertible. In this example they were sampled from 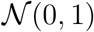. Finally the connectivity matrix of the network **M** was computed from **M** = **UΛU**^-1^.

The simulated activity of the network is shown in Figure 3a (top left). The initial condition was specified to satisfy Equation 5. The bottom left of Figure 3a shows the network activity rescales along the time axis according to the peak times of each neuron. It is evident that the activations of the neurons are rescaled version of each other, confirming that the constraints derived above indeed lead to scale-invariant sequential activity.

**Figure 3:**
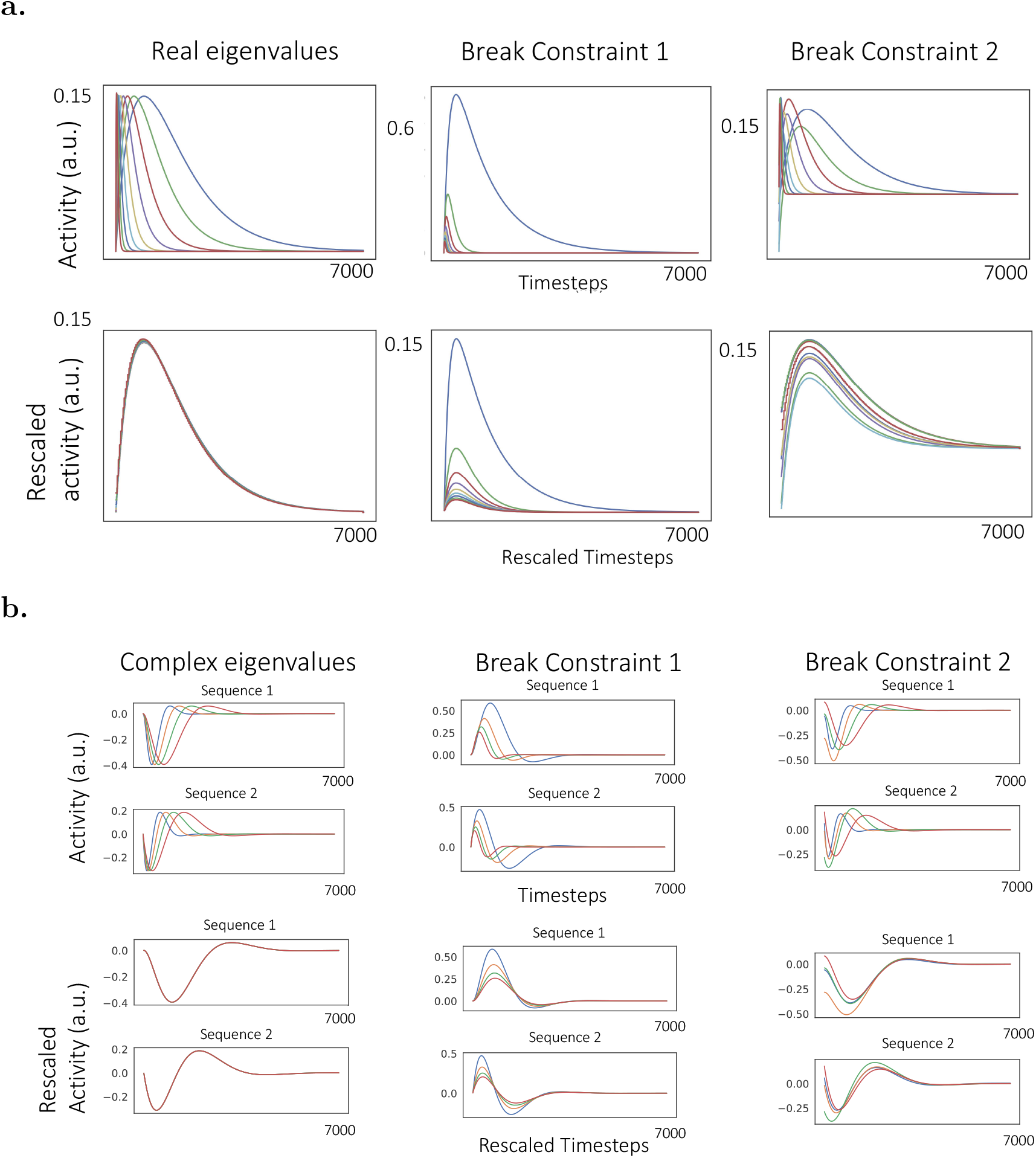
Generating scale-invariant neural sequences with simple (a) and complicated (b) single neuron dynamics. Using the analytical result, we constructed networks with specific connectivity matrices so that they generate scale-invariant sequential activity (left column). We also broke each of the two constraints derived in Section 2 and showed that the resulting network activity breaks scale-invariance (middle and right columns) **a.** The network with real eigenvalues gives rise to simple unimodal single cell temporal receptive fields (**a**, left top, each line represents the activity of one neuron in the network). The activity of different neurons overlap with each other when rescaled according to their peak times (left bottom). The network activity becomes not scale-invariant when Constraint 1 was broken by choosing linearly spaced eigenvalues or Constraint 2 was broken by adding noise to the motifs (**a**, middle and right columns, see text for details). **b.** The network with complex eigenvalues gives rise to two sequences of neuronal responses with more complex temporal dynamics (**b** left top, see text and Appendix A.2 for details). The responses of cells in each sequence are rescaled version of each other in time (**b** left bottom). When each of the two constraints is broken, the network activity becomes not scale-invariant (right two columns).

We also broke each of the two constraints above and showed that the resulting activity became no longer scale-invariant (Figure 3a, right two panels). To break Constraint 1 (geometrically spaced eigenvalues), linearly spaced eigenvalues in the same range were used in constructing the matrix **Λ** instead of geometrically spaced eigenvalues. To break Constraint 2 (translation-invariant eigenvectors), a random vector was added to each row of the matrix **U** where each entry was sampled from 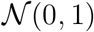, making each motif different. As a result, both manipulations generated activity that was no longer scale-invariant (Figure 3a, bottom right two panels).

### 3.2 Complicated single-cell tuning curves

Temporal coding needs not result in a simple unimodal sequential code. Instead, it could also be embedded in the collective activity of neuronal populations where single neurons may exhibit highly complex dynamics (e.g. Machens et al., 2010). In this subsection we show that the framework we presented above is sufficiently rich to allow for more complex single cell dynamics. In the previous example, the connectivity matrix has real eigenvalues. Therefore it can only give rise to single cell dynamics that are linear combinations of exponential functions. On the other hand, in this subsection we will show that more complicated temporal dynamics can be generated if the connectivity matrix has complex eigenvalues.

Following a similar procedure as detailed in Section 3.1, the connectivity matrix with N = 20 neurons was set up so that all of its eigenvalues were complex, and their real and imaginary parts both formed geometric progressions with the same real common ratio (see Appendix A.2 for details). As mentioned in Section 2.2, 2.3 and in more detail in Appendix A.1, the eigenvalues and eigenvectors form complex conjugate pairs, and the network activity contains two distinct scale-invariant sequences. The simulated and rescaled activity are shown in Figure 3b (left). The neural activity exhibits more complicated dynamics but at the same time maintains scale-invariance for each sequence.

We also broke each of the two constraints using similar protocols as in the previous example. To break Constraint 1, eigenvalues with linearly spaced real and imaginary parts in the same range were used instead of geometrically spaced ones. To break Constraint 2, multiplicative noise sampled from 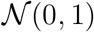 was used, so that each matrix element *U_ij_* becomes *U_ij_* (1 + *ϵ_ij_*) where each *ϵ_ij_* was sampled from 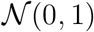. As shown in Figure 3b (right two panels), the resulting neural activity is no longer scale-invariant.

### 3.3 A special case: Laplace and inverse Laplace transforms

It should be noted that the geometric progression of time constants required by Constraint 1 does not necessarily have to be an emergent property of the network, but can instead be driven by physiological properties of single cells (Loewenstein and Sompolinsky, 2003; Fransén et al., 2006; Tiganj et al., 2015; Liu et al., 2019). Consequently, scale-invariant sequential activity could also be generated by feedforward networks where the neurons in the first layer receive inputs and decay exponentially with a spectrum of geometrically-spaced intrinsic time constants, and the neurons in the second layer are driven by the first layer via translation-invariant synaptic weights, implementing the eigendecomposition in Equation 3 explicitly.

One such model has been proposed by Shankar and Howard (Shankar and Howard, 2013). It is a two-layer feedforward neural network. In that model, the first layer neurons **F** encode the Laplace transform of the input and have exponentially decaying firing rates with a spectrum of decay constants.

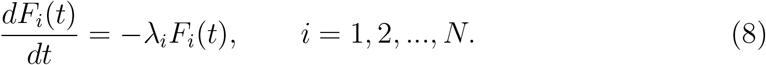

The activity of the first layer constitutes a scale-invariant sequential activity (see Equation 2). It is also the basis functions that make up any general scale-invariant sequential activity (see Equation 3).

To generate sequential neural activity, the neurons in the second layer 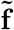 compute the inverse Laplace transform of the first layer under the Post approximation (Post, 1930).^2^

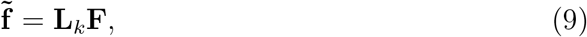

where **F** is the activity vector of all the first layer neurons and 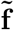 is that of the second layer neurons. The matrix **L**_*k*_ is a discretized approximation of the inverse Laplace transform of the *k*th order (Shankar and Howard, 2013).

From Equation 8 and Equation 9, the feedfoward dynamics above is equivalent to linear recurrent dynamics involving only the second layer neurons 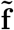:

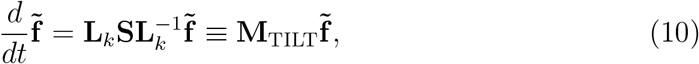

where **S** is a diagonal matrix consisting of the single cell time constants λ_*i*_’s in Equation 8.

Therefore the dynamics of the neurons in the second layer are equivalent to the one generated by a linear recurrent network with connectivity matrix **M**_TILT_. Because the matrix representation of the inverse Laplace transform **L**_*k*_ is approximated by taking derivatives of nearby nodes, it has the same motif across columns (for details see Shankar and Howard, 2013). Therefore, the model in Shankar and Howard (2012), although a feedforward network, is a special case in the family of linear recurrent networks that can generate scale-invariant sequential activity. It might be the case that that the exponentially decaying basis functions are indeed maintained by a separate population of neurons, and the downstream neurons consititute a “dual” population. Such exponentially decaying cells have recently been identified in lateral entorhinal cortex (Tsao et al., 2018; Bright et al., 2019), whose downstream regions have been identified as locations for time cells (Pastalkova et al., 2008; MacDonald et al., 2011; Kraus et al., 2013; Salz et al., 2016).

## 4 Comparison with common network models

Scale-invariance puts stringent constraints on the architecture of recurrent neural networks. To illustrate this, in this section we consider two widely-used neural network models that do not generate scale-invariant sequential activity. Section 4.1 considers a simple chaining model; the following section considers a random network. We will see that the connectivity matrices for these two widely-used models violate the constraints derived above in Section 2 and that these models do not support scale-invariant sequential activity. We will compare these two networks with the two scale-invariant example networks described previously. All the networks in this section are simulated with *N* = 20 neurons.

### 4.1 Simple Feedforward Chaining Model

The simplest possible model to generate sequential activity is a simple feedforward chaining model, which will be analyzed in this subsection. The connectivity matrix of the studied model is non-diagonalizable, therefore the constraints derived above in Section 2.2 and 2.3 do not apply exactly. However, as discussed in Appendix A.4, the *distinct* eigenvalues should still form a geometric progression even when the connectivity is non-diagonalizable to allow scale-invariant activity. We will demonstrate that a chaining model composed of elements with the same time constant cannot meet this requirement for scale-invariant sequential activity.

In this simple model the activity of the *k*th neuron in a chain of *N* neurons obeys

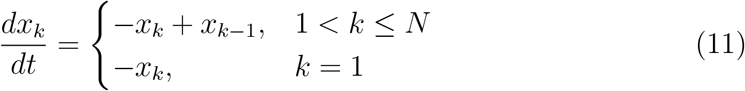

Note that all of these neurons have the same time constant which is here set to 1. Because an activation in the first unit spreads gradually across the network, this model generates sequential activity. Each neuron does not respond instantaneously to its input but has some finite integration time resulting in a spread of activity across time. As the activation spreads across the chain, the spread in time accumulates. However, as can be shown via the Central Limit theorem, this sequential activity is not scale-invariant because the peak time of the *k*th unit goes up linearly in *k* but the width of the peak goes up with 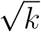 (Liu et al., 2019). Here we show that the eigenspectrum of the connectivity matrix of this simple model (Eq. 11) does not satisfy the constraints derived from Section 2.

Figure 4a (left) shows the connectivity matrix of the chaining model described in Eq. 11. This connectivity matrix has a simple motif that repeats across rows; the self-interaction term gives a —1 on the diagonal and the chaining term gives a +1 on the off-diagonal. This connectivity matrix is non-diagonalizable and has a single degenerate eigenvalue of −1 (Figure 4a, middle left). According to Appendix A.4, the distinct eigenvalues of a non-diagonalizable connectivity matrix should still follow a geometric progression for the network to allow scale-invariant activity. The connectivity matrix of the chaining model violates this condition. Therefore, although the network generates sequential activity (Figure 4a, middle right), the activity of different neurons are not rescaled versions of each other (Figure 4a, right).

**Figure 4:**
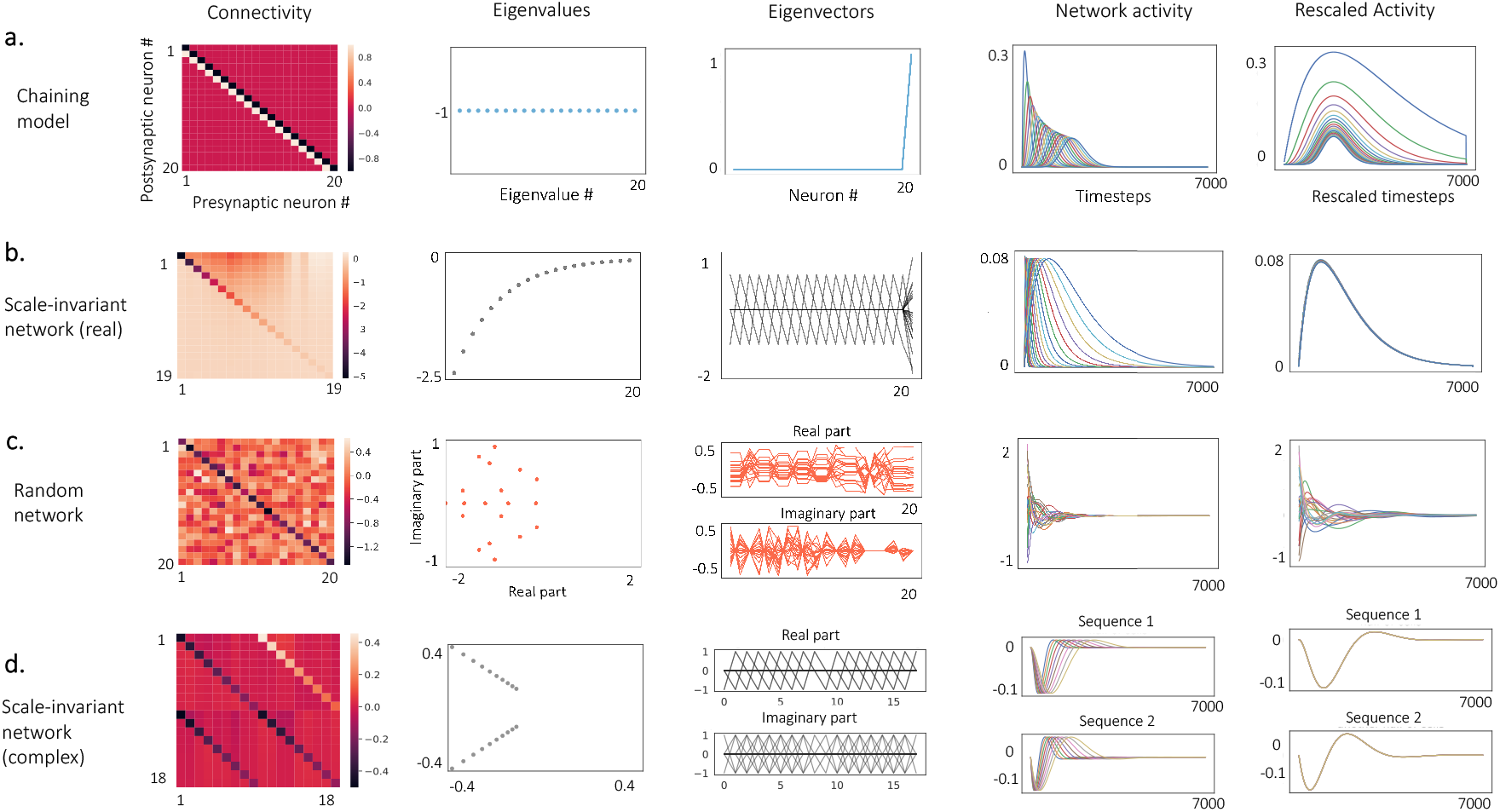
Comparison with common network models. **a, b.** Comparison between the simple feedforward model (**a**) and a network constructed from Section 3.1 that generates scale-invariant sequential activity (scale-invariant network [real], **b**). For the simple feedforward chaining model, its connectivity matrix is non-diagonalizable. Because the eigenvalues of its connectivity matrix are not geometrically spaced (**a**, middle left), the network activity does not have the scaleinvariant property (**a**, right two panels). In contrast, the scale-invariant network has eigenvalues that are geometrically spaced (**b**, middle left) and eigenvectors that consist of the same motif (**b**, middle, aside from the boundary effect), resulting in the connectivity that consists of excitation from neurons earlier in the sequence to the ones later in the sequence and inhibition in the opposite direction (**b**, left, only neurons in the sequence are shown. Same below). Consequently, it allows scaleinvariant sequential activity (**b** right two panels). **c, d.** Same comparison between an instance of a random recurrent network (**c**) and a network constructed from Section 3.2 (scale-invariant network [complex], **d**). The eigenvalues of the connectivity of a random network are not geometrically spaced (**c**, middle left). The eigenvectors do not consist of the same motif (**c**, middle). Consequently its activity is not scaleinvariant (**c**, right two panels). In contrast, the connectivity of the scale-invariant network has two geometric progression of eigenvalues that are complex conjugate with each other (**d**, middle left). Its eigenvectors also contain complex conjugate pairs, each with a repeating motif (**d**, middle). Consequently the scale-invariant network can generate two scale-invariant sequences (**d**, right two panels).

For contrast, Figure 4b illustrates the connectivity matrix, eigenvalues and eigenvectors for the scale-invariant network with real eigenvalues described previously in Section 3.1. In illustrating the connectivity matrix, we have ordered the neurons according to the order in which they are activated and we have only included the neurons in the sequence (the same for the complex example below). Recall that this matrix was constructed to obey the two constraints and has already been shown to generate scale-invariant sequential activity. Consequently, by construction, the eigenvalues are geometrically spaced and the eigenvectors are translated versions of one another (Figure 4b, middle left and middle panels). Although the connectivity matrix clearly has a rich structure, the rows of the connectivity matrix are certainly not translated versions of one another. The entries above the diagonal tend to be more negative, indicating that the connections from neurons later in the sequence to the neurons earlier in the sequence tend to be more inhibitory than the connections in the opposite direction (Figure 4b, leftmost panel). It is not at all obvious why this specific connectivity matrix yields scale-invariant sequential activity. This is much more clear from examining the eigenvalues and eigenvectors.

### 4.2 Random recurrent networks

Nonlinear neural networks with random connectivity matrices are able to generate chaotic activity (Sompolinsky et al., 1988). With appropriately trained weights, they are also able to generate sequential neural activity similar to that obtained in actual recordings (Rajan et al., 2016). However, generic linear random neural networks without training cannot produce scale-invariant, sequential activity due to the conflict with the two constraints derived above in Section 2.

Random neural networks have eigenvalue spectrums that are uniformly distributed inside a unit disc in the complex plane (Rajan and Abbott, 2006; Girko, 1985), therefore not geometrically spaced as required by scale-invariance (Section 2.2). We simulated the activity of an instance of a linear network with a random connectivity matrix and computed its eigenvalues and eigenvectors (Figure 4c). The network dynamics is described by

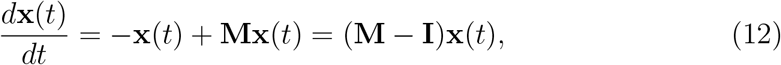

where each element in **M** was sampled from a Gaussian distribution with mean 0 and variance 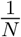.

The connectivity matrix **M** — **I** is shown in Figure 4c (left). The eigenvalues of the connectivity matrix are approximately uniformly distributed in a unit disc centered at (−1,0) (Figure 4c, middle left), confirming the results from Rajan and Abbott (2006) and Girko (1985). The eigenvectors also apparently do not have any translation-invariant structure (Figure 4c, middle). Consequently, the network activity does not have the scale-invariant property (Figure 4c, right two panels). In contrast, the network constructed in the same way as in Section 3.2 has two geometric progressions of complex eigenvalues (Figure 4d, middle left) that are complex conjugates of each other, and each half of the eigenvectors consist of the same motif (Figure 4d, middle). Consequently, it allows two sequences of scale-invariant activity (Figure 4d, right two panels).

## 5 Discussion

In this paper we study the constraints on the connectivity of a linear network for it to generate scale-invariant sequential neural activity. It is analytically shown that two conditions need to hold for the structure of the connectivity matrix when it has real, distinct eigenvalues. First of all, it must have geometrically-spaced eigenvalues. Second, its eigenvectors must contain the same motif with a translation-invariant structure. Intuitively, the same motif in different eigenvectors pick out different timescales by the same proportion, and the geometric spacing of timescales further ensures the activity of different cells are rescaled with each other. The generated activity is highly dynamic, providing a possibility for the dynamic coding of working memory (Stokes, 2015). It was also shown that a straightforward generalization leads to the constraints for networks with complex eigenvalues (for details see Appendix A.1). Analogous but more complicated constraints hold for networks with degenerate eigenvalues (Appendices A.3 and A.4).

### 5.1 Plausibility for a geometric progression of timescales

Geometrically spaced network eigenvalues can generically emerge from multiplicative cellular processes (Amir et al., 2012). Mechanistically, a spectrum of timescales could be generated by positive feedback in recurrent circuits. It could also be an inherent property of single neurons (Loewenstein and Sompolinsky, 2003; Franséen et al., 2006; Tiganj et al., 2015; Liu et al., 2019). *In vitro* slice experiments and modeling works have shown that a spectrum of slow timescales can be obtained in single cells by utilizing the slow dynamics of calcium-dependent currents (Loewenstein and Sompolinsky, 2003; Egorov et al., 2002; Fransén et al., 2006; Mongillo et al., 2008; Tiganj et al., 2015; Liu et al., 2019). *In vivo* experiments also showed a spectrum of timescales in cortical dynamics. Bernacchia et al. (2011) showed that in monkey prefrontal, cingulate and parietal cortex, reward modulates neural activity multiplicatively with a spectrum of time constants (Bernacchia et al., 2011).

### 5.2 Weber-Fechner Law

The constraints for diagonalizable matrices with real unique eigenvalues suggest a deep connection to the Weber-Fechner law. The Weber-Fechner law relates internal psychophysical scales to external physical variables. A quantity is said to obey the Weber-Fechner law if a psychological quantity *p* is related to a physical quantity *S* as *p* = *k* log *S* for an appropriate choice of units. The Weber-Fechner law is perhaps the oldest quantitative relationship in psychology (Fechner, 1912) and holds at least approximately for a wide range of physical variables.

Consider a set of receptors responding to some physical variable *S* with a receptive field center s¿. If the receptive field centers follow a geometric progression, a logarithmic mapping between receptor number and *S* naturally results. Logarithmic spacing of receptive fields seems to be a common property of receptive fields in sensory systems (Merzenich et al., 1973; Van Essen et al., 1984; Schwartz, 1977). However, analogous logarithmic spacing is also observed for receptive fields over non-sensory variables such as numerosity (Nieder and Miller, 2003; Nieder and Dehaene, 2009). The receptive fields of time cells in multiple brain regions are certainly compressed (Mello et al., 2015; Kraus et al., 2013; Tiganj et al., 2018), but it has not been quantitatively established that this compression is logarithmic (Howard, 2018).

The constraints for diagonalizable matrices with unique real eigenvalues developed here are closely analogous to the requirements for a Weber-Fechner scale. The geometric progression of eigenvalues places the eigenvalues on a logarithmic scale such that the *n*th eigenvalue goes up exponentially with n. The requirement that the eigenvectors are translated versions of the same motif is analogous to an translation-invariant tiling along some dimension. Thus, scale-invariance in linear recurrent networks can be interpreted as a translation-invariant tiling of receptors along a logarithmic scale.

### 5.3 The eigenvectors of scale-invariant networks are localized

The second constraint we derived requires that connectivity matrices with real, distinct eigenvalues should have eigenvectors that consist of the same motif localized at different entries. It is hard to imagine how this kind of translation-invariant eigenvectors could arise generically in neural circuits. However the eigenvectors that satisfy this constraint are a special case of “localized eigenvectors”, which have been studied extensively first in condensed matter physics and later in theoretical neuroscience. Anderson first argued that the eigenvectors of matrices whose elements are random and concentrated on the diagonal are exponentially localized (Anderson, P., 1958). Later numerical and analytical studies confirmed that localized eigenvectors indeed arise in neural networks in the presence of a gradient in the strength of the local interactions (Chaudhuri et al., 2014) or global inhibition, as in the case of ring attractor networks (Tanaka and Nelson, 2019). However, it should be pointed out that the procedures described in this paper do not necessarily generate local interactions where the elements *M_ij_* decays with |*i* — *j*|, as can be seen in Figure 4b. Furthermore, our constraint on the eigenvectors is more stringent than only requiring them to be localized: the different localized “patches” need to be the same as well.

### 5.4 Scale-invariant sequence gives logarithmic growth in cumulative dimensionality

In a recent study, Cueva et al. (2019) showed that the cumulative dimensionality of the population activity during working memory increases with a decreasing speed (Cueva et al., 2019). This is consistent with the network activity generated by linear networks that satisfy the two constraints we derived. Translation-invariant eigenvectors ensure that each eigenmode contributes one unique dimension to the activity. Geometrically spaced eigenvalues ensure that one eigenmode would be suppressed per unit time on a logarithmic scale. Therefore the cumulative dimensionality of scale-invariant sequential activity would increase with the logarithm of time. Furthermore, notice that any affine transformation on the neural trajectory would not change the cumulative dimensionality. Therefore the same relationship for cumulative dimensionality would hold even if the eigenvectors are not translation-invariant. A geometric progression of eigenvalues is sufficient to generate linear dynamics whose cumulative dimensionality increases with the logarithm of time.

### 5.5 Chaining models

Goldman (2009) proposed a class of linear network models that are able to generate sequential activity. In these networks, the feedforward dynamics are constructed by building up feedforward interactions between orthogonal Schur modes and are hidden in the collective network dynamics (Goldman, 2009). Thanks to the hidden feedforward dynamics, the networks can sustain its activity far longer than the timescale constrained by the eigenvalue spectrum of the network. The example studied in Section 4.1 is the simplest model proposed in Goldman (2009). Although this simple model does not allow scale-invariant sequential activity, this does not argue that any network constructed in the way in Goldman (2009) cannot generate scale-invariant activity.

### 5.6 Non-normal networks

Because the eigenvetors in a scale-invariant network are translated versions of each other, they do not form an orthonormal basis. Therefore, the networks that generate scale-invariant sequential activity naturally belong to the family of non-normal networks (White et al., 2004; Ganguli et al., 2008; Goldman, 2009). Non-normal networks have been shown to have many desired properties such as extensive memory capacity where the number of timesteps over which the network state retains information about the past stimuli scales linearly with the size of the network (White et al., 2004; Ganguli et al., 2008). The current work shows that scale-invariance could be another potential computational benefit for non-normal networks.

### 5.7 Locally-interacting integral transform networks

Shankar studied a model where the activity of individual nodes can be effectively described by a filtered input through a set of scale-invariant kernel functions with different timescales (Shankar, 2015). It was shown that if this transformation were to be implemented by a network involving only local (possibly non-linear) interactions between nodes with similar timescales, the forms of the kernel functions are strongly constrained: they can only be given by a linear combination of the inverse Laplace transforms (c.f. Section 3.3). In this work we consider a slightly different problem: the interactions between different nodes are constrained to be linear, but non-local interactions are also allowed since the connectivity matrix is not constrained to be local. In this case, the activity given by the inverse Laplace transform (Equation 10) covers a subspace of all possible solutions. It is still notable that these two related approaches converge on similar results.

## Acknowledgements

This work was supported by ONR MURI award N00014-16-1-2832, ONR DURIP award N00014-17-1-2304 and NIBIB R01EB022864.

## A Appendix

The constraints derived in Section 2 are the sufficient and necessary conditions for generating scale-invariant activity in a linear recurrent network, but only when the network connectivity matrix has real, distinct eigenvalues. In this series of appendices we discuss the modifications to the constraints when the connectivity matrix does not meet this requirement. We start out by discussing matrices with complex, distinct eigenvalues (Appendix A.1). The results from this appendix have already been stated in the main text. We then use these results to describe the setup of the example network in Section 3.2 in Appendix A.2. The case when the network connectivity has degenerate eigenvalues and is diagonalizable is discussed in Appendix A.3. The case when it is non-diagonalizable is discussed in Appendix A.4. Finally in Appendix A.5 we discuss the sensitivity of the scale-invariant property to perturbations in the initial condition and the network connectivity.

### A.1 Connectivity matrices with complex eigenvalues

We consider the case when all the eigenvalues are complex. It will be become obvious below that a mixture of real and complex eigenvalues will make the network activity further deviate from being strictly scale-invariant (c.f. Equation 2), therefore we do not consider this case in this paper.

Since network connectivity matrices are real, their complex eigenvalues and eigenvectors come in conjugate pairs. Consequently, the two constraints in Section 2 should be stated in a slightly different way. The activity of cell *i* is still

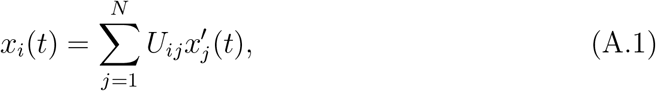

where the modes are

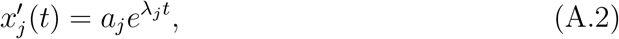

except that the eigenvalues λ_*i*_ are now complex. Therefore the derivation in Section 2 carries over. However, since the eigenvectors form complex conjugate pairs, the matrix **U** in the diagonalization procedure **M** = **UΛU**^-1^ should be of the following form:

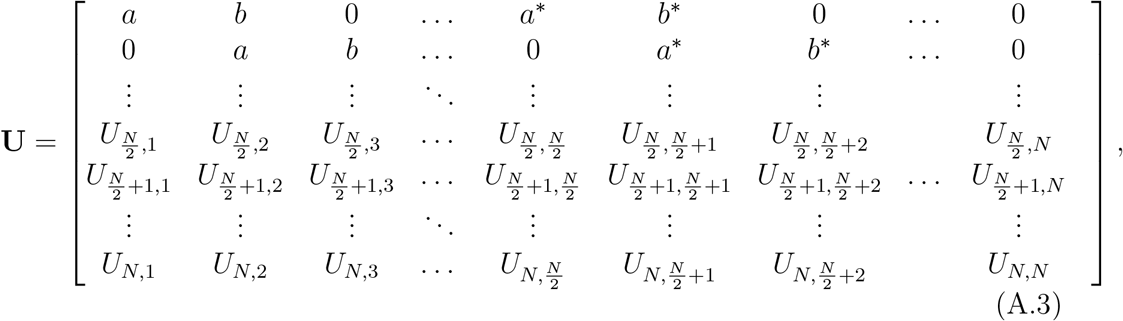

where the motif is (*a,b*). Notice that the lower half of the matrix is not determined. This is due to the constraint that there has to be a complex conjugate for every eigenvector, and therefore the motif can only be translated 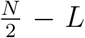 times, rather than *N — L* times as in the case of real eigenvalues (c.f. Section 2.3). The bottom half of the matrix cannot simply repeat the upper half as this will make **U** not invertible. Therefore, it is not possible to satisfy the scale-invariant condition in the main text that any pair of responses are rescaled version of each other (c.f. Equation 2). Instead, the network can only have half of its cells be scale-invariant (cells 1 to 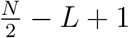).

But we can do a little better if we allow the bottom half of the matrix to repeat the structure of the upper half, only with a different motif:

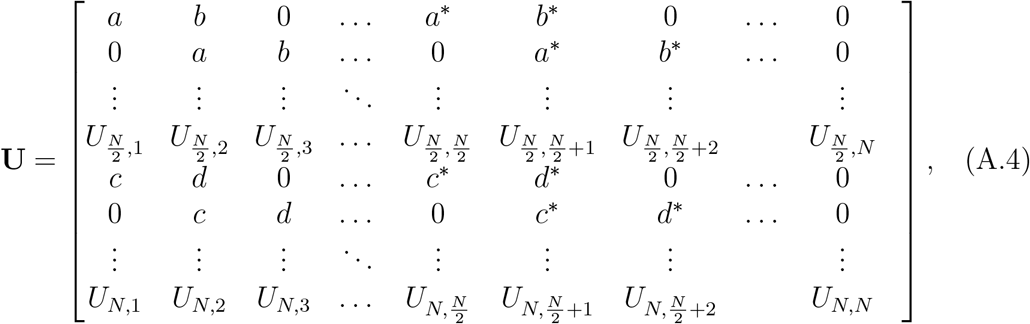

This way, the network will generate two different scale-invariant sequences. Cells within each sequence has responses that are rescaled version of each other, but cells in different sequences are not rescaled version of each other. This is the way the example network in Section 3.2 was constructed.

To summarize, linear recurrent networks with connectivity matrices whose eigenvalues are all complex cannot generate scale-invariant activity in the strict sense as defined by Equation 2. The closest scenario is to have two different scale-invariant sequences, each composed of half of the cells in the network. In this case, the constraint on the eigenvalues is that they form two geometric progressions that are complex conjugate with each other, with the same real common ratio for both the real and imaginary parts. The constraint on the eigenvectors is that each of the eigenvector should either be a translated version of another eigenvector or its complex conjugate. We applied these results to construct the example network in Section 3.2, as detailed in the next appendix.

Finally, it is not hard to see that if there is a mixture of real and complex eigenvalues, the network can at best generate three distinct scale-invariant sequences. Since this scenario is even further from the strict definition of scale-invariance (c.f. Equation 2), we do not consider this case in this paper.

### A.2 The setup of the example network with complex eigenvalues

For the example in Section 3.2, the real and imaginary parts of the eigenvalues were both chosen to be geometrically spaced between −0.15 and −0.48 (−0.15, −0.17, …, −0.48), and the complex conjugates were also included. Therefore, the matrix **Λ** in the diagonalization procedure **M** = **UΛU**^-1^ is

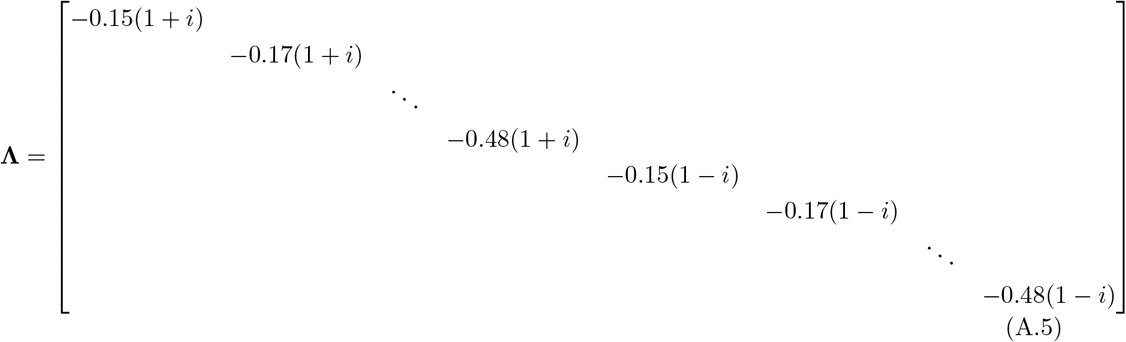

To satisfy the constraint that the eigenvectors form complex conjugate pairs, we constructed the motif so that the matrix **U** is in the form of Equation A.4 with two repeating motifs, where *a* =1 + *i, b* = –1 – *i, c* =1 — *i, d* = —1 + *i*. In other words the matrix **U** is

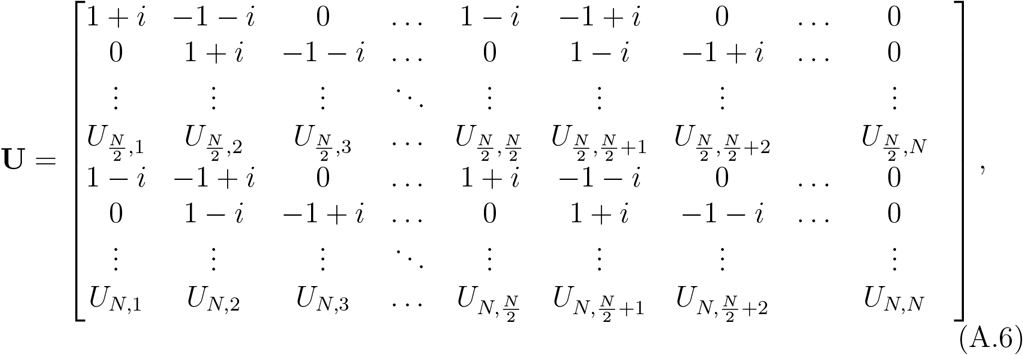

where the unspecified elements are chosen randomly from a uniform distribution from [0,11.

### A.3 Diagonalizable connectivity matrices with degenerate eigenvalues

A matrix with degenerate eigenvalues could be diagonalizable or non-diagonalizable (defective). The modifications to the constraints are different for these two cases. In this appendix we discuss these modifications when the network connectivity is diagonalizable and in the next appendix we focus on non-diagonalizable matrices. In both cases, degeneracy in the eigenvalues decreases the number of distinct responses in the scale-invariant sequence because this number is equal to the number of *distinct* eigenvalues of the connectivity. In both this and the next appendices, we assume that the eigenvalues of the connectivity matrix are all real. It is straightforward to generalize the results to the case of complex eigenvalues based on the discussions in Appendix A.1.

For diagonalizale matrices with degenerate eigenvalues, the activity of cell i is still

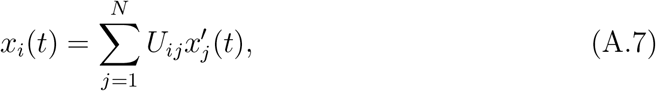

where the modes are

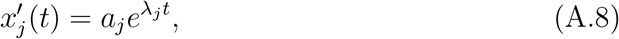

except that some of the eigenvalues λ_*j*_ could be the same. Therefore, for scale-invariance to hold, the modes that contribute to the summation in Equation A.7 for any pair of cells should have eigenvalues that are a multiple of each other (e.g. for cell 1, the eigenvalues that contribute to the summation in Equation A.7 are {−1, −1, −2, −4}, then for cell 2 they could be {−2, −2, −4, −8}, etc.). A necessary condition for this to hold is that the *distinct* eigenvalues of the connectivity matrix form a geometric progression. Therefore, the constraint on the eigenvalues is as follows:

> For a linear recurrent network whose connectivity matrix is diagonalizable with degenerate, real eigenvalues to generate scale-invariant activity, the *distinct* eigenvalues must form a geometric progression (c.f. Section 2.2).

The constraint on the eigenvectors is more complicated due to the degeneracy of eigenvalues. We here offer a compact statement for a sufficient condition on the eigenvectors: During the diagonalization of the matrix **M** = **UΛU**^-1^, if we rearrange the eigenvalues in the diagonal form of the connectivity **Λ** such that they form blocks of distinct eigenvalues (e.g. λ_1_ = — 1,λ_2_ = — 2,λ3 = — 4, λ_4_ = — 1,λ_5_ = —2, etc.), then at the block level, the matrix **U** would satisfy the original constraint that its columns (i.e. the eigenvectors) consist of the same motif (c.f. Section 2.2) at translated entries. In general, the eigenvectors would at least to be localized.

### A.4 Non-diagonalizable connectivity matrices

A non-diagonalizable (or defective) matrix can be transformed into a block-diagonal matrix called its Jordan normal form by a similarity transformation. We will show that when the connectivity matrix is non-diagonalizable, its *distinct* eigenvalues still need to form a geometric progression. However, as for diagonalizable matrices with degenerate eigenvalues (Appendix A.3), the constraint on the eigenvectors is more complicated. In what follows, we assume all eigenvalues are real. It is straighforward to generalize the results to the case of complex eigenvalues based on the discussions in Appendix A.1.

To reach this conclusion, we first review some basic results about the Jordan normal form. A non-diagonalizable connectivity matrix **M** can be transformed into a block-diagonal matrix via a similarity transformation: **M** = **U**^-1^ **JU**. The columns of **U** are called the generalized eigenvectors of **M** and **J** is called the Jordan normal form of **M. J** is a block-diagonal matrix where the diagonal elements of each block is an eigenvalue λ of the matrix **M**. It also has off-diagonal elements 1 to the right of each diagonal element, i.e. *J*_*i,i*+1_ = 1, except for the last row of each block. Consequently, it is not hard to see that the activity of each cell is a linear combination of modes

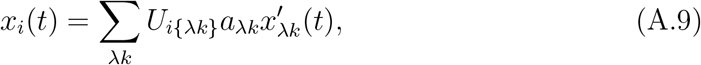

where we use 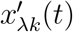 to represent the mode corresponding to the *k*th last row of the block with eigenvalue λ. Here, the modes will not all be exponential functions, but a product of a polynomial and an exponential

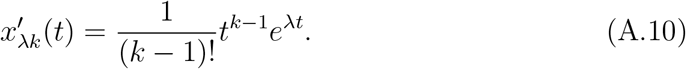

Based on Equations A.9 and A.10 above, it is quite straightforward to come up with the constraints on scale-invariance in the case of non-diagonalizable matrices. For any pair of cells, the modes that contribute to their activity should have series of eigenvalues λ that are a multiple of each other as well as the same *sequence* of *k*. For example, if there are two modes contributing to the activity of cell 1, with (*k*_1_, λ_1_) = (1, –1) and (*k*_2_, λ_2_) = (2, —2), then the two contributing modes for cell 2 could have (*k*_1_,λ_1_) = (1, —2) and (*k*_2_, λ_2_) = (2, —4), and similar relationships hold for any pair of cells. As a special case, for diagonalizable connectivity matrices, the contributing modes for any cell have *k* = 1 for all λ. The necessary condition for the contributing eigenvalues for different cells to be a multiple of each other is that the distinct eigenvalues of the connectivity form a geometric progression. Therefore, the constraint on the eigenvalues is as follows:

> For a linear recurrent network whose connectivity matrix is non-diagonalizable and has real eigenvalues to generate scale-invariant activity, the *distinct* eigenvalues of the connectivity matrix should form a geometric progression (c.f. Section 2.2).

However, the constraint on the eigenvectors is harder to state in a compact manner because of the degeneracy of the eigenvalues. The best we can say is that each eigenvector would have to be localized, i.e. the range of the non-zero elements would have to be narrow.

### A.5 Robustness to noise

In this appendix we show that deviations from the constraints derived in Section 2 causes a graceful degradation in scale-invariance. We performed numerical experiments where we perturbed the connectivity matrix of the network in Section 3.1 using Gaussian multiplicative noise: 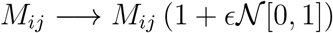 and the initial condition with Gaussian additive noise: 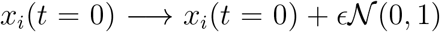. We chose an additive noise for the initial condition because a multiplicative noise would only rescale the initial state for the example in Section 3.1. The degree of scale-invariance was measured in terms of the ratio of the peak times between successive cells in the sequence. If the sequence is scale-invariant, this ratio should equal to the common ratio of the geometric progression of eigenvalues for all pairs of successive cells.

Figure A.1a and d show the distributions of the ratio obtained by 100 realizations of the perturbation on the connectivity and the initial condition, respectively. As can be seen, the distributions become more spread out around the noiseless values as the noise amplitudes *ϵ* are increased. Figure A.1b and e show that the deviation from scale-invariance as measured by the average difference from the noiseless ratio increases linearly with the noise amplitude *ϵ*. Figure A.1c and f show the raw and rescaled activity for two realizations of the perturbation with different noise levels. These results show that the degree of scale-invariance decreases linearly with the level of noise in both the initial condition and the connectivity matrix. Generally, a small perturbation to a matrix changes its eigenvalues and eigenvectors by an amount linear in the perturbation, and a small deviation in the initial condition of a linear dynamical system causes the subsequent trajectory to diverge from the original trajectory by an amount linear in the deviation. Therefore it is expected that the deviation from scale-invariance increases linearly with noise strength.

**Figure A.1:**
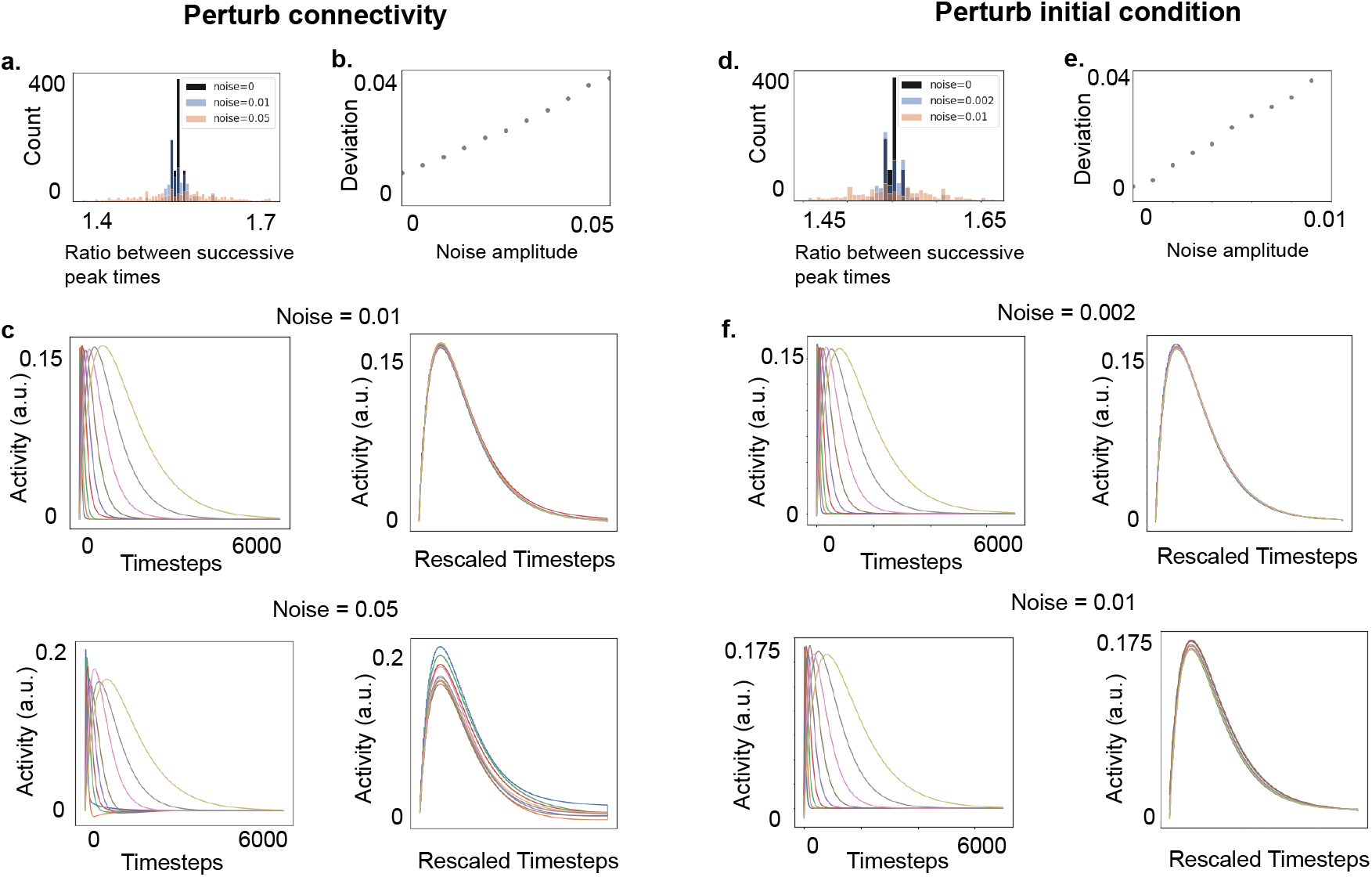
Degree of scale-invariance decreases gracefully with noise. We perturbed the connectivity and the initial condition with Gaussian multiplicative and additive noise respectively for 100 independent runs (see texts for details) and measured the deviation from scale-invariance by the spread of the distribution of the ratio between peak times for successive cells in the sequence. For the perturbation on the connectivity matrix, the distribution of the ratio is more spread out for larger noise (**a**). The mean difference from the noiseless ratio increases linearly with the noise amplitude (**b**). **c.** The raw (left) and rescaled (right) activity for two example runs with different noise levels are shown. **d-f.** The same results for the perturbation on the initial condition. Since in both cases the deviation from scale-invariance is linear in noise strength, scale-invariance does not depend on fine-tuning the connectivity matrix or the initial condition.

## Footnotes

1 When there is a mixture of real and complex eigenvalues, there will be at least 3 sequences, which is further away from the strict definition of scale-invariance in Equation 2. Therefore we do not consider this case in this paper.

2 Note that this transformation is written as 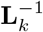 in other papers.

